# Intranasal dantrolene nanoparticles inhibit inflammatory pyroptosis in 5XFAD mice brains

**DOI:** 10.1101/2024.11.25.625293

**Authors:** Piplu Bhuiyan, Wenjia Zhang, Rebecca Chae, Kyulee Kim, Lauren St. Louis, Ying Wang, Ge Liang, Huafeng Wei

## Abstract

**Background:** This study investigates the effects of intranasal dantrolene nanoparticles on inflammation and programmed cell death by pyroptosis in 5XFAD Alzheimer’s Disease (AD) mice.

**Methods:** 5XFAD and wild type (WT) B6SJLF1/J mice were treated with intranasal dantrolene nanoparticles (5 mg/kg), daily, Monday to Friday, for 12 weeks continuously, starting at 9 months of age. Blood and brain were harvested at 13 months of age, one month after completion of 12 weeks intranasal dantrolene nanoparticle treatment. Blood biomarkers function of liver (Alanine transaminase, ALT), kidney (Creatinine), and thyroid (TSH: Thyroid-stimulating hormone) were measured using ELISA. The changes of whole brain tissue proteins on Ca^2+^ release channels on membrane of endoplasmic reticulum (type 2 ryanodine and type 1 InsP3 receptors, RyR-2 and InsP3R-1), lipid peroxidation byproduct malondialdehyde (MDA)-modified proteins, 4-HNE, pyroptosis regulatory proteins (NLR family pyrin domain containing 3 (NLRP3), cleaved caspase-1, full length or N-terminal of Gasdermin D (GSDMD), cytotoxic (IL-1, IL-18, IL-6, TNF-a) and cytoprotective (IL-10) cytokines, astrogliosis (GFAP), microgliosis (IBA-1) and synapse proteins (PSD-95, Synapsin-1) were determined using immunoblotting. Body weights were monitored regularly.

**Results:** Intranasal dantrolene nanoparticles significantly inhibited the increase of RyR-2 and InsP3R-1 proteins, MDA-modified proteins, 4-NHE, pyroptosis regulatory proteins (NLRP3, cleaved caspase-1, N-terminal GSDMD), cytotoxic cytokine (IL-1β, IL-18, IL-6, TNF-α), biomarkers for astrogliosis (GFAP) and microgliosis (IBA-1), and the decrease of cytoprotective cytokine (IL-10) and synaptic proteins (PSD-95, synpasin-1). Intranasal dantrolene nanoparticles for 12 weeks did not affect blood biomarkers for function of liver, kidney, and thyroid, not did it change body weight significantly.

**Conclusion:** Intranasal dantrolene nanoparticles significantly inhibit the increase of RyR-2 and InsP_3_R-1 Ca^2+^ channel receptor proteins, ameliorate activation of the pyroptosis pathway and pathological inflammation, and the associated loss of synapse proteins. Intranasal dantrolene nanoparticles for three months did not affect liver, kidney and thyroid functions or cause other side effects.

## Introduction

Although mechanisms unclear, increasing evidence suggest that disruption of intracellular Ca^2+^ homeostasis and associated pathologies including inflammation and synapse dysfunction play critical role of pathologies in Alzheimer’s disease (AD)^1^. The abnormally increased glutamate and associated excitotoxicity via overactivation of N-methyl-D-aspartate (NMDA) receptors (NMDAR) result in the overactivation of ryanodine (RyRs) and InsP_3_ (InsP_3_R) receptors located on membrane of endoplasmic reticulum (ER) and associated excessive Ca^2+^ release from ER, leading to depletion of ER Ca^2+^ and pathological elevation of cytosol and mitochondrial Ca^2+^ concentrations, detrimental to cell survival^2, 3^. The type 2 RyRs (RyR-2) is pathologically increased in brains of AD patients^4^. The numbers and activity of RyRs pathologically increase AD in preclinical studies^5, 6^. Similarly, the numbers and activity of InsP_3_R also pathologically increased in AD^7, 8^. The upstream Ca^2+^ dysregulation results in downstream mitochondrial dysfunction, oxidative stresses, initiation of pyroptosis pathway and pathological inflammation, eventually leading to synapse and cell/neurons damage^9^, and depression and cognitive dysfunction^10, 11^. A drug with potency to inhibit inflammation and programmed cell death by inflammatory pyroptosis is expected to be neuroprotective in AD.

Dantrolene, a RyRs antagonist, is an FDA approved drug for treatment of malignant hyperthermia, muscle spasm, and neuroleptic syndrome with tolerable side effects and occasional liver toxicity at high doses^12^. Dantrolene inhibits the common upstream critical Ca^2+^ dysregulation and has been shown to be neuroprotective against many neurodegenerative diseases, including cerebral ischemia, Huntington’s disease, spinocerebellar ataxia, ALS and seizures^9, 13^. We and others have demonstrated that dantrolene abolishes or ameliorates the cognitive dysfunction in multiple AD animal models^11, 14-16^. Our recent studies pioneered a novel approach of administrating intranasal dantrolene nanoparticles to increase its brain/blood concentration ratio, promotes dantrolene CNS therapeutic effects and minimizes its peripheral side effects/organ toxicity^11, 17, 18^. Intranasal dantrolene nanoparticles, as a disease-modifying drug, significantly inhibited memory loss with no side effects or muscle or liver toxicity after 10 months treatment in 5XFAD mice^11^.

In this study, we investigated the effects of intranasal dantrolene nanoparticles chronic treatments on inflammation and inflammatory pyroptosis in 5XFAD transgenic mice. Our results demonstrated that intranasal dantrolene nanoparticles administered for 12 consecutive weeks from 9 to 12 months old significantly inhibited inflammation and inflammatory pyroptosis in 5XFAD mice brains, illustrating a new mechanism of dantrolene neuroprotection in AD.

## Materials and Methods

### Animals

All procedures were approved by the Institutional Animal Care and Use Committee (IACUC) at the University of Pennsylvania. Four pairs of 5XFAD mice (B6SJL-Tg (APPSwFlL on, PSEN1^*^M146L^*^L286V) 6799Vas/Mmjax) and wild type mice (B6SJLF1/J) were purchased from the Jackson Laboratory (Bar Harbor, ME) and were bred. These 5XFAD transgenic mice over-express mutant human APP with the Swedish (K670N, M671L), Florida (I716V), and London (V717I) Familial Alzheimer’s Disease (FAD) mutations along with human PS1 harboring two FAD mutations, M146L and L286V. Food and water were available in the cage. All mice were weaned no later than one month of age and were genetically identified by polymerase chain reaction (PCR) analysis before weaning. At this time, mice were divided into different cages according to age and gender, with no more than five mice per cage. Both male and female mice were used in this study.

### Dantrolene administration

Dantrolene (Sigma, St Louis, MO) was dissolved in the Ryanodex Formulation Vehicle (RFV: 125 mg mannitol, 25 mg polysorbate 80, 4mg povidone K12 in 5mL of sterile water and pH adjusted to 10.3) to form crystalline nanoparticles19, similarly as in our previous publications^11, 17^. For intranasal administration, the final concentration of dantrolene was 5 mg/mL as we described previously, the mice were held and fixed by the scruff of their necks with one hand and with the other hand given a total of 1μL/gram of body weight of dantrolene nanoparticle solution or RFV. A mouse weighing 20 g would be given 20μL solution. The solution was slowly delivered directly into the mouse’s nose. Care was taken to make sure that mice were minimally stressed, and that the solution stayed in the nasal cavity and did not enter the stomach or lungs. Both WT and 5XFAD mice at 9 months old were treated with intranasal dantrolene nanoparticles (5 mg/kg), daily, from Monday to Friday, for 12 continuous weeks.

### Euthanasia and tissue collection

As we described previously^11, 18^ animals were deeply anesthetized with 2–4% isoflurane delivered through a nose cone, and the concentration was adjusted according to the animals’ response to a toe pinch. The animal’s skin was prepped, and an incision made to open the chest and expose the heart. Blood was collected for the serum study from the heart using a syringe equipped with a 27G needle. The blood was centrifuged at 1400 rpm at 4°C for 30 min, the supernatant collected and frozen at –80°C. The animals were euthanized by trans cardiac perfusion and exsanguination with cold phosphate-buffered saline. Then, the brain was dissected and stored in –80°C for western blot.

### Immunoblotting

Total brain tissues were extracted by homogenization using cold RIPA buffer (#9806S, Cell Technology, USA) supplemented with protease inhibitor cocktails (P8340 Roche). The brain homogenates were rocked at 4°C for 90 minutes and then centrifuged at 14,000 rpm (Brushless Microcentrifuge, Denville 260D) for 20 minutes at 4°C to remove cell debris. After collecting the supernatant, the protein concentration was measured using a BCA protein assay kit (Pierce, Rockford, IL 61101 USA). Briefly, equal amounts of protein (50µg/lane) were loaded onto 4-20% gel electrophoresis of mini-protein TGX precast (Cat. #4561094, BIO-RAD) and transferred to polyvinylidene difluoride (PVDF) membranes (Immobilon-P, MERK Millipore, Ireland) using wet electrotransfer system (BIO-RAD, USA). Following the transfer membrane blocking with 5% BSA (Sigma-Aldrich) for 1 hour, then the PVDF membrane was incubated with primary antibodies overnight at 4 °C including RYR2, InsP_3_R1, MDA-modified protein, 4-HNE, NLRP3, Human procaspase-1/ cleaved caspase-1 p20, GSDMD, Cleaved GSDMD, IL-1β, IL-18, IL-6, TNF-α, GFAP, IL-10, PSD95, Synapsin-1, IBA-1, GAPDH [Details antibodies information table 1]. After that, the membranes were incubated with a secondary antibody including anti-Mouse IgG1 HRP-linked; Anti-rabbit IgG, HRP-linked that was conjugated to horseradish peroxidase and washed with Tris-buffered saline containing 0.2% Tween-20 (TBST). Subsequently, TBST was used for washing the membranes three time for 10 minutes. After incubation with secondary antibodies, proteins band were visualized using ECL Western Blotting Detection Reagents (Cytiva, Amersham, UK) and quantified for band intensity using ImageJ software (National Institutes of health, Bethesda, MD, USA).

**Table 1:** Antibodies applied in immunoblotting.

|  | Cat. and Manufacture | Dilution |
| --- | --- | --- |
| **Primary antibody** |  |  |
| Anti- RYR2 | 19765-1-AP; Proteintech | 1:1000 |
| Anti- InsP3R1 | # PA1-901, Invitrogen | 1:1000 |
| Anti-MDA-modified protein | MA5-27560, Invitrogen | 1:1000 |
| Anti-4 Hydroxynonenal | (ab46545), Abcam | 1:1000 |
| Anti- NLRP3 | 68102-1-Ig; Proteintech | 1:2000 |
| Anti-Human procaspase-1/<br>cleaved caspase-1 p20 | #2225S, Cell signaling Technology, USA | 1:1000 |
| Anti-GSDMD | ab155233; Abcam | 1:1000 |
| Anti- Cleaved GSDMD | #36425S; Cell signaling Technology, USA | 1:1000 |
| Anti-IL-1β | 31202S, Cell signaling Technology, USA | 1:1000 |
| Anti-IL-18 | 10663-1-AP; Proteintech | 1:3000 |
| Anti-IL-6 | # M620; Invitrogen | 1:1000 |
| Anti-TNF-α | 60291-1-Ig; Proteintech | 1:2000 |
| Anti-GFAP | #3670; Cell signaling Technology, USA | 1:1000 |
| Anti-Iba1 | # 17198; Cell signaling Technology, USA | 1:1000 |
| Anti-IL-10 | #12163S, Cell signaling Technology, USA | 1:1000 |
| Anti-PSD95 | #2507; Cell signaling Technology, USA | 1:1000 |
| Synapsin-1 | #5297; Cell signaling Technology, USA |  |
| GAPDH | MA5-15738, Thermo Fisher Scientific | 1:10,000 |
| **Secondary antibody** |  |  |
| Anti-Mouse IgG1; HRP -linked | # PA1-74421, Thermo Fisher Scientific | 1:10,000 |
| Anti-rabbit IgG, HRP-linked | #7074; Cell signaling Technology, USA | 1:1000 |

### Serum alanine aminotransferase (ALT) activity

Serum ALT activity, an indicator of liver function, was measured using an Alanine Aminotransferase (ALT) Activity Colorimetric Assay Kit (ab105134, ABCAM, Inc, Waltham, MA, USA), a method of ELISA assay. According to the manufacturer’s instructions. Briefly, 20μl serum was diluted in a total 100μl reaction mixture, including 86μl ALT Assay Buffer, 2μl OxiRed Probe, 2μl ALT Enzyme Mix, and 10μl ALT Substrate, to analyze the pyruvate transformed from a-ketoglutarate with alanine. A standard curve was generated at the same time, using pyruvate concentrations of 0, 2, 4, 6, 8, and 10 nmol/well. The optical density (OD) at 570 nm was measured at 10 min (A1) then again at 60 min (A2) after incubating the reaction at 37°C. The pyruvate concentration was measured in a linear range of the standard curve. ALT activity was calculated using the formulation: ALT activity = (A2-A1)/ (50^*^0.02) ^*^1 mU/ml.

### Serum creatinine (Cr) activity

Serum creatinine, an indicator of kidney function, was measured using a Creatinine Colorimetric Assay Kit (ab65340, Abcam, Inc, Waltham, MA, USA) according to the manufacturer’s instructions. We measured the serum creatinine for the 5XFAD mice. Briefly, 50μl serum was diluted in a total 100μl reaction mixture, including 42μl Cr Assay Buffer, 4μl Creatinase, 2μl Cr Enzyme Mix, and 2μl Creatinine Probe. A standard curve was generated, using creatinine concentrations of 0, 2, 4, 6, 8, and 10 nmol/well. The optical density (OD) at 570 nm was measured at 60 min after incubating the reaction at 37°C. The creatinine concentration was measured in a linear range of the standard curve. The trendline equation (Sa) was calculated based on the standard curve data. The creatinine concentration was calculated using the formulation: creatinine concentration =Sa/50 nmol/μl.

### Serum TSH (Thyroid Stimulating Hormone) activity

Serum TSH, a hormone that indicated the thyroid gland function, was measured using a TSH Colorimetric Assay Kit (CHM01J578, Thomas scientific, Inc, Swedesboro, NJ, USA) according to the manufacturer’s instructions. We measured the serum thyroid stimulating hormone for the 5XFAD mice. Briefly, 10μl serum was diluted in a total 50μl with 40μl of sample diluent and was incubated for 30 min at 37°C. After 5 times of washing with washing buffer, 50μl HRP-Conjugate reagent was added. After 30 min incubation at 37°C and five times washing, the reaction was stopped with stop solution. The optical density (OD) at 450 nm was measured within 15 min. At the same time, a standard curve was generated, using TSH concentrations of 0, 50,100,200,400,600 and 900 pg/ml. The sample concentration was calculated by using straight line regression equation of the standard curve.

### Statistical Analysis

All data were represented as mean ± 95% confidence index. Statistical analyses were employed with GraphPad Prism (Version 10.3.1, CA, USA). Comparisons of more than two groups were conducted by two-way ANOVA with Tukey’s multiple comparison test (MCT). P<0.05 was considered statistically significant.

## Results

### Dantrolene inhibited the pathological increase of type 2 ryanodine receptor (RYR-2) and type 1 InsP_3_R (InsP_3_R-1) in 5XFAD mice brains

In aged mice at 13 months old, RyRs-2 were pathologically and dramatically increased in 5XFAD mice, which was robustly inhibited by intranasal dantrolene nanoparticles for 12 weeks between 9 and 12 months old (Fig. 1A, C). As InsP_3_R-1 activity also significantly increased in AD cells^8^ or animal models^7^, we also determined the effects of dantrolene treatment on changes of InsP_3_R-1 proteins using immunoblotting. Similarly, intranasal dantrolene nanoparticles treatments for 12 weeks robustly inhibited the pathological increase of InsP_3_R-1 protein in 5XFAD mice brain (Fig. 2B, D).

**Figure 1:**
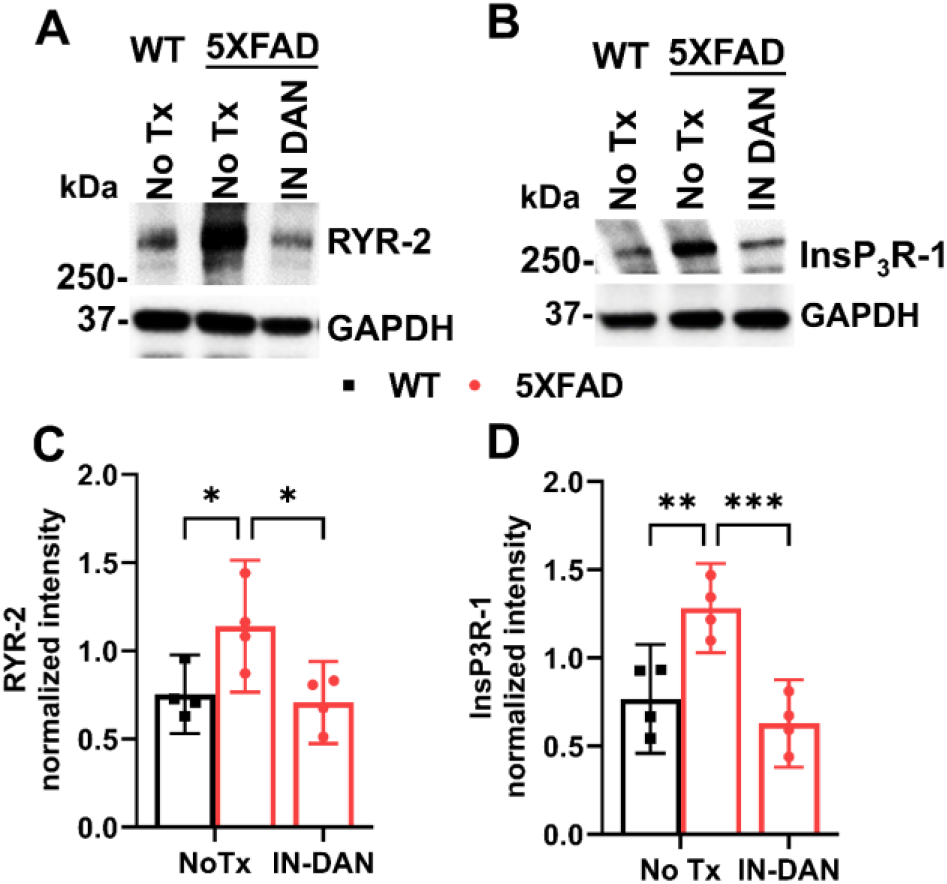
Dantrolene inhibits the pathological increase of type 2 ryanodine receptor (RyR-2) and type 1 InsP_3_R (InsP_3_R-1) proteins in 5XFAD mice brains. Wild type (WT) or 5XFAD mice at 9 months old were treated with intranasal dantrolene nanoparticles (IN-DAN, 5 mg/kg), daily, Monday to Friday for 12 consecutive weeks. Mice brains were harvested at 13 months old. Immunoblot was used to determine the protein levels of RyR2 **(A, C)** and InsP_3_R-1 **(B, D)**. GAPDH was used as the housekeeping protein. Data are means with 95% confidence interval (Cl) from 4 separate mice in each experimental group (N=4) and were analyzed using the two-way ANOVA followed by Tukey’s multiple comparison test. ^**, ***^, P<0.01, P<0.001.

**Figure 2.**
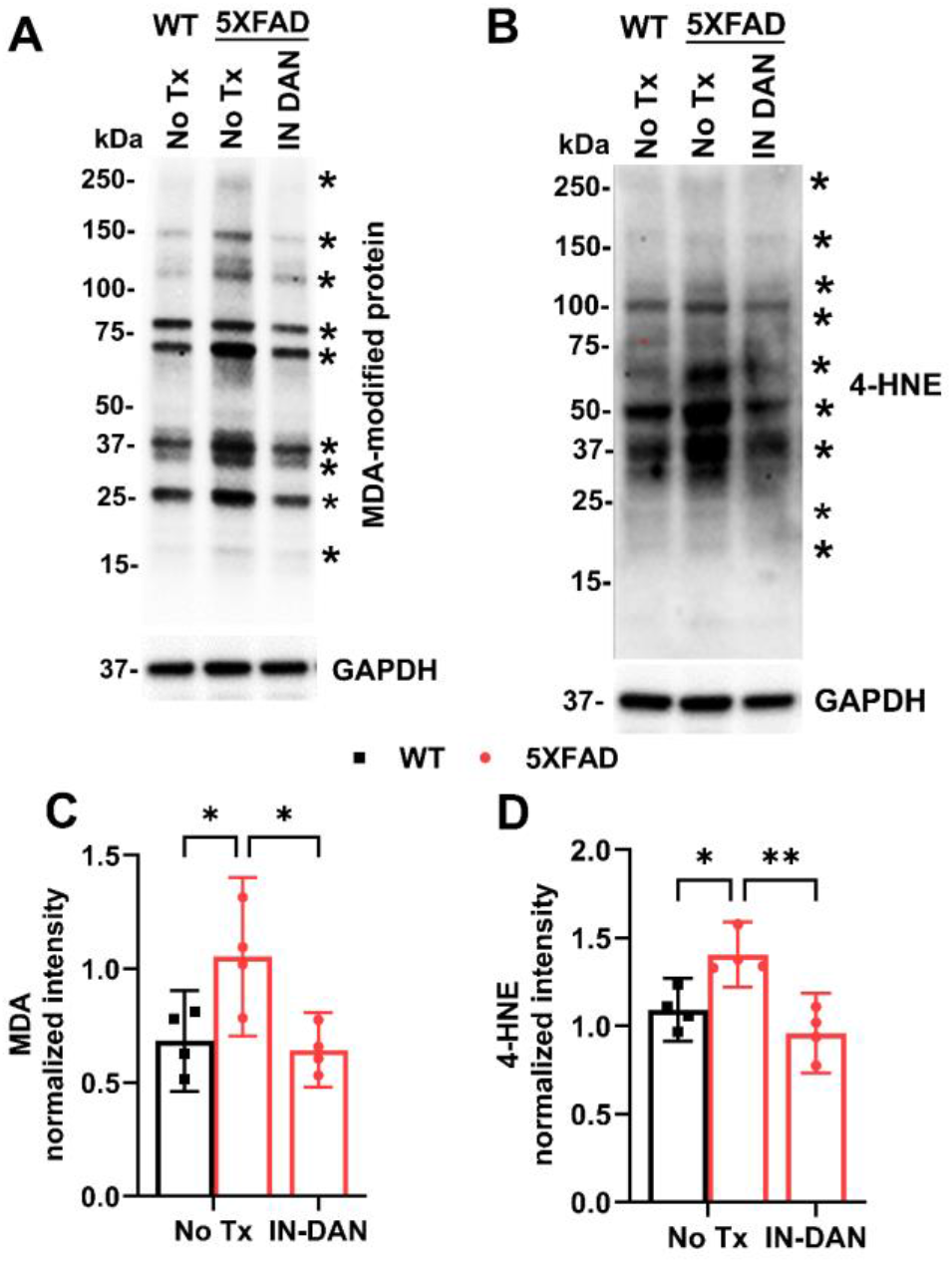
Dantrolene inhibits oxidative stress in 5XFAD mice brains. Wild type (WT) or 5XFAD mice at 9 months old were treated with intranasal dantrolene nanoparticles (IN-DAN, 5 mg/kg), daily, Monday to Friday for 12 consecutive weeks. Mice brains were harvested at 13 months old. Representative Western blot was used to measure malondialdehyde (MDA, lipid peroxidation byproduct) modified proteins (**A, C)** and 4-HNE **(B, D)** and quantified all the modified proteins marked by **\***. Data represented as means with 95% confidence interval (Cl) from 4 separate mice brain proteins (N=4) in each experimental group and were analyzed using the two-way ANOVA followed by Tukey’s multiple comparison test. ^*, **^, P<0.05, P<0.01.

### Dantrolene suppressed the oxidative stress in 5XFAD mice

We measured protein oxidation by determining oxidized biomolecules by-products, malondialdehyde (MDA) modified proteins and 4-Hydroxynonenal (4-HNE). MDA modified proteins (Fig. 2A, C) and 4-HNE (Fig. 2B, D) were significantly increased in the 13 months old 5XFAD mice brain, which could be robustly inhibited by intranasal dantrolene nanoparticle treatment for 12 weeks from 9 to 12 months of age.

### Dantrolene inhibits pathological activation of the pyroptosis pathway in 5XFAD mice brains

We determined the levels of primary regulatory proteins of pyroptosis, NLR family pyrin domain containing 3 (NLRP3), caspase-1, Gasdermin D full length (GSDMD-FT) and N terminal Gasdermin D (N-GSDMD) in wild type versus 5XFAD mice. The core pyroptosis initiation protein NLRP3 significantly increased in 13-month-old 5XFAD mice brains compared with wild type control, which could be robustly inhibited by the intranasal dantrolene nanoparticles treatment for 12 consecutive weeks (Fig. 3A, B). Consistently from the same mice brain proteins, intranasal dantrolene nanoparticles treatment robustly inhibited downstream pathological elevation of caspase-1 (Fig. 3C, D). Dantrolene treatment did not change full length Gasdermin D (GSDMD-FL, Fig 3E, F), but robustly inhibited the elevation of N-terminal Gasdermin D (N-GSDMD, Fig. 3G, H). In addition, the pyroptosis related cytotoxic cytokines IL-1β (Fig. I, J) and IL-18 (Fig. 3K, L) were significantly increased in 5XFAD mice brains, which were robustly inhibited by intranasal dantrolene nanoparticles treatment.

**Figure 3.**
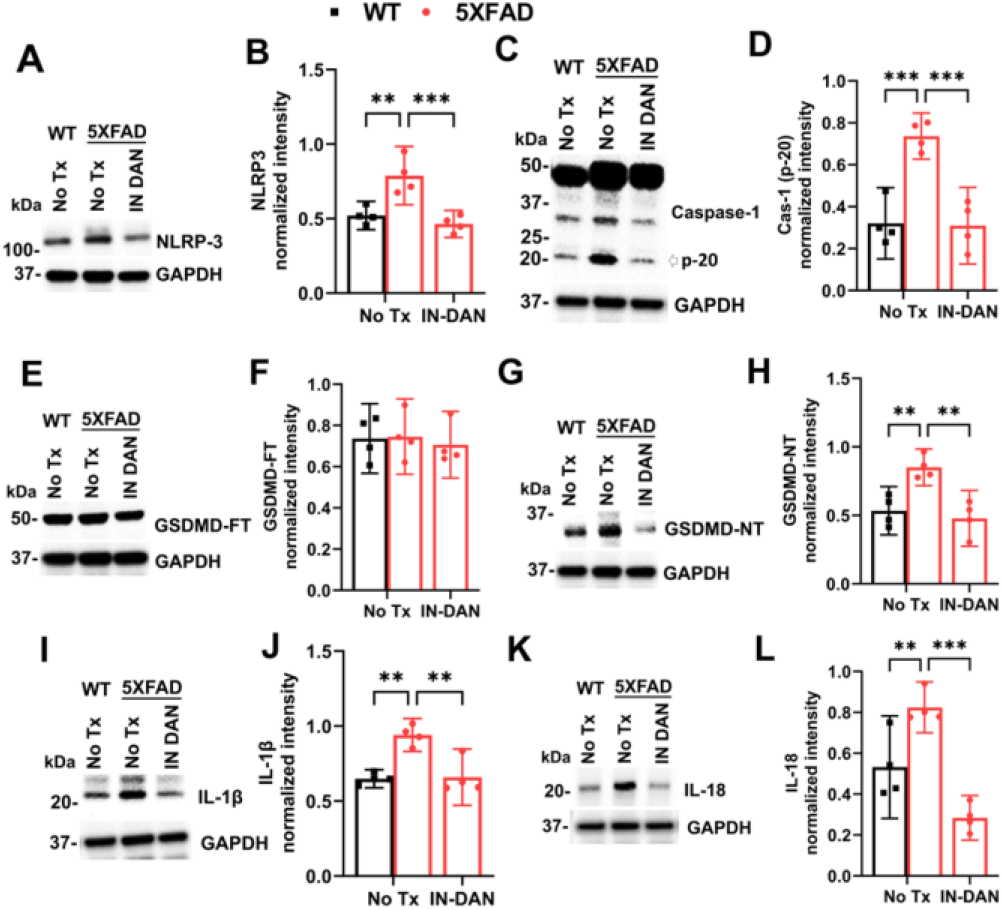
Dantrolene inhibits pathological activation of the pyroptosis pathway in 5XFAD mice. Wild type (WT) or 5XFAD mice at 9 months old were treated with intranasal dantrolene nanoparticles (IN-DAN, 5 mg/kg), daily, Monday to Friday for 12 consecutive weeks. Mice brains were harvested at 13 months old. Representative Western blots and associated statistical analysis were used to determine changes of critical regulatory proteins of pyroptosis activation pathway (**A and B**. NLR family pyrin domain containing 3 (NLRP3); **C and D**. Caspase-1; **E and F**. Gasdermin D full length (GSDMD-FL); **G and H**. Gasdermin D N terminal (GSDMD-NT); **I and J**. IL-1β; and **K and L**. IL-18). GAPDH was used as the loading control. Data are means with 95% confidence interval (Cl) from 4 separate mice brains (N=4) in each experimental group and were analyzed using two-way ANOVA followed by Tukey’s multiple comparison test. ^**, ***^ P<0.01, P<0.001.

### Dantrolene inhibits the pathological elevation of neurotoxic cytokines but promotes neuroprotective cytokines in 5xFAD mice brains

Besides the cytotoxic cytokines related to inflammatory pyroptosis (IL-1β, Fig. 3I, J, IL-18, Fig. 3K, L), we also examined other inflammation related cytotoxic cytokines. IL-6 was significantly increased in 5XFAD mice brains (Fig. 4A, D), so was the TNF-a (Fig. 4B, E), while both could be robustly inhibited by treatment with intranasal dantrolene nanoparticles. On the other hand, intranasal dantrolene nanoparticles significantly increased neuroprotective cytokines IL-10 in 5XFAD mice (Figure 4C, F).

**Figure 4:**
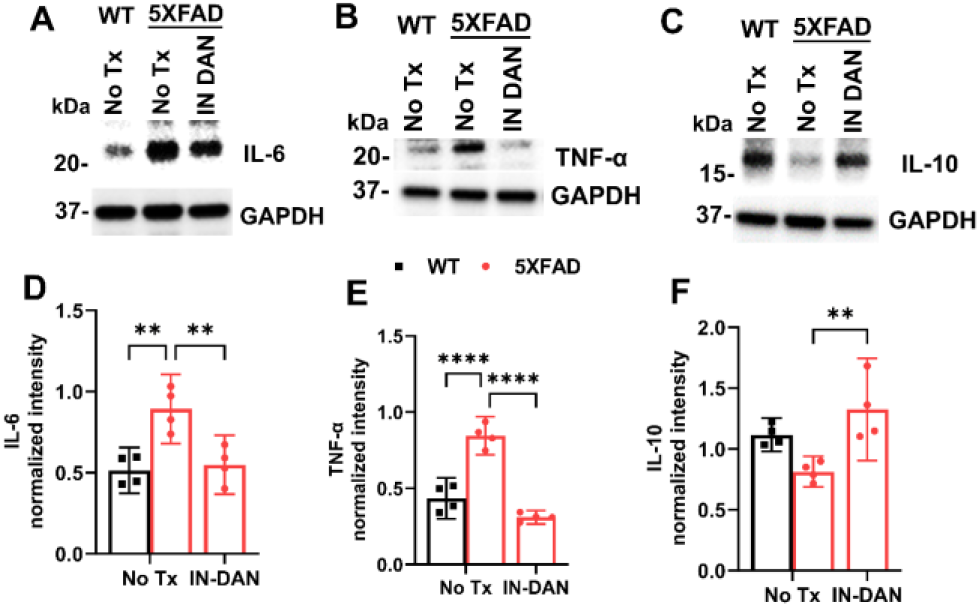
Dantrolene inhibits the pathological increase of neurotoxic cytokines but increase neuroprotective cytokines in 5XFAD mice. Wild type (WT) or 5XFAD mice at 9 months old were treated with intranasal dantrolene nanoparticles (INDAN, 5 mg/kg) nanoparticles, daily, Monday to Friday for 12 consecutive weeks. Mice brains were then harvested at the age of 13 months old. Representative Western blot and associated statistical analysis of neurotoxic cytokines proteins IL-6 (**A, D**), TNF-α (**B, E**) and neuroprotective cytokine IL-10 (**C, F)** were determined and analyzed. GAPDH as loading control. Data are means with 95% confidence interval (Cl) from 4 separate mice brains (N=4) and were analyzed using two-way ANOVA followed by Tukey’s multiple comparison test. ^**, ****^, P<0.01, P<0.0001.

### Dantrolene inhibits astrogliosis and microgliosis in 5XFAD mice brains

Both astrogliosis and microgliosis contribute to pathological neuroinflammation^20^. GFAP, a biomarker protein for astrogliosis, was significantly increased in 5XFAD mice brain compared to WT controls, which was robustly inhibited by intranasal dantrolene treatments (Fig. 5A, C). Similarly, the biomarker protein for microgliosis, IBA-1, was significantly elevated in 5XFAD mice brains compared to WT controls, which was also robustly inhibited by intranasal dantrolene treatments (Fig. 5B, D).

**Figure 5.**
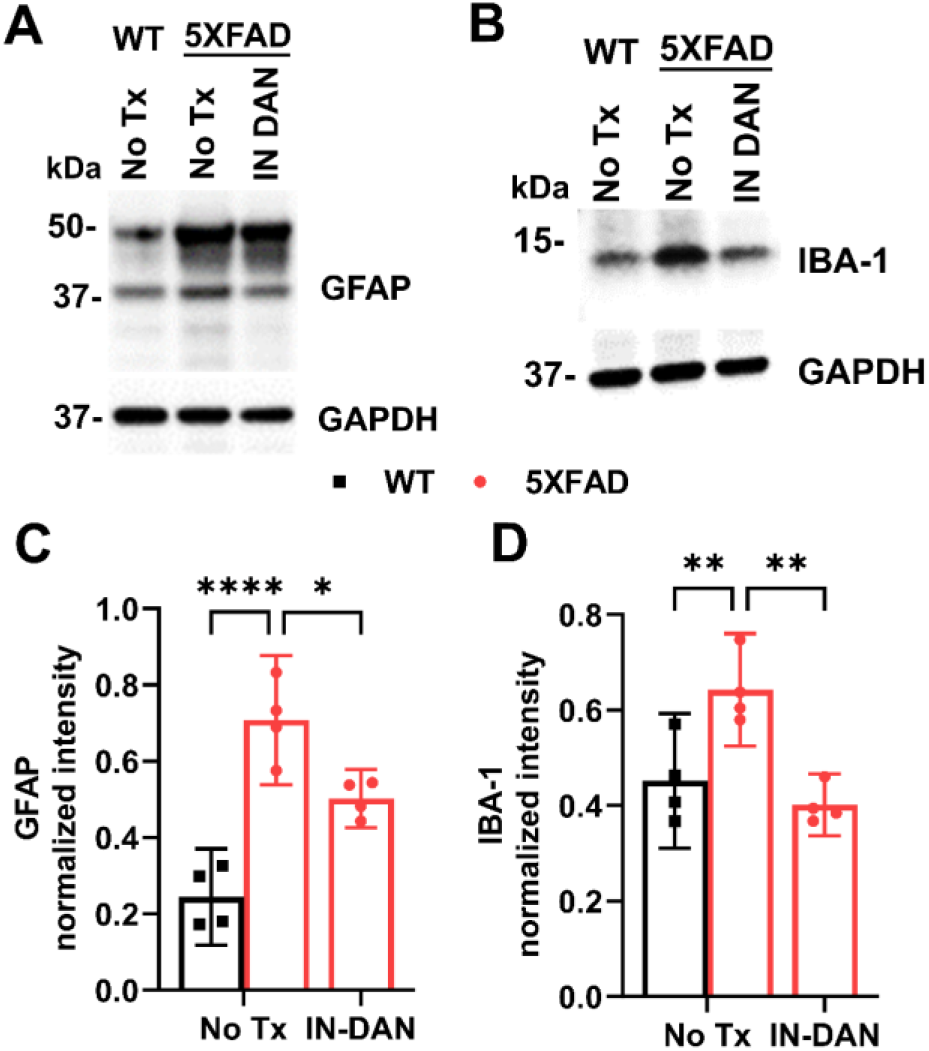
Dantrolene inhibited astrogliosis/microgliosis in 5XFAD mice brains. Wild type (WT) or 5XFAD mice at 9-monthold were treated with intranasal dantrolene nanoparticles (IN-DAN, 5 mg/kg) nanoparticles, 5 days/week for 3 consecutive months. Mice brains harvested at 13 months old. Immunoblot was used to determine protein biomarkers of astrogliosis (GFAP) **(A, C)** and microgliosis (IBA-1) **(B, D)**. Data are means with 95% confidence interval (Cl) from 4 separate mice brains (N=4) and analyzed using two-way ANOVA followed by Tukey’s multiple comparison test. ^*, **, ****^ P<0.05, P<0.01, P<0.0001, respectively.

### Dantrolene inhibited the decrease of synaptic proteins in 5XFAD mice

Our previous study showed that the use of dantrolene resulted in a significant inhibition of synaptic proteins loss and impaired synaptogenesis in prefrontal cortical neurons that were derived from the iPSCs of Alzheimer’s disease patients^5^. Therefore, we determined the effects of intranasal dantrolene nanoparticles on synapse proteins levels in aged 5XFAD mice. PSD-95 protein levels significantly decreased 5XFAD mice brains at 13 months of age, which was robustly inhibited by the intranasal dantrolene nanoparticles treatments for 12 weeks from 9 to 12 months old (Fig. 6A, C). Furthermore, dantrolene treatment also robustly inhibited the decrease of synpasin-1, another important synapse protein, in these 5XFAD mice brains (Fig. 6B, D).

**Figure 6:**
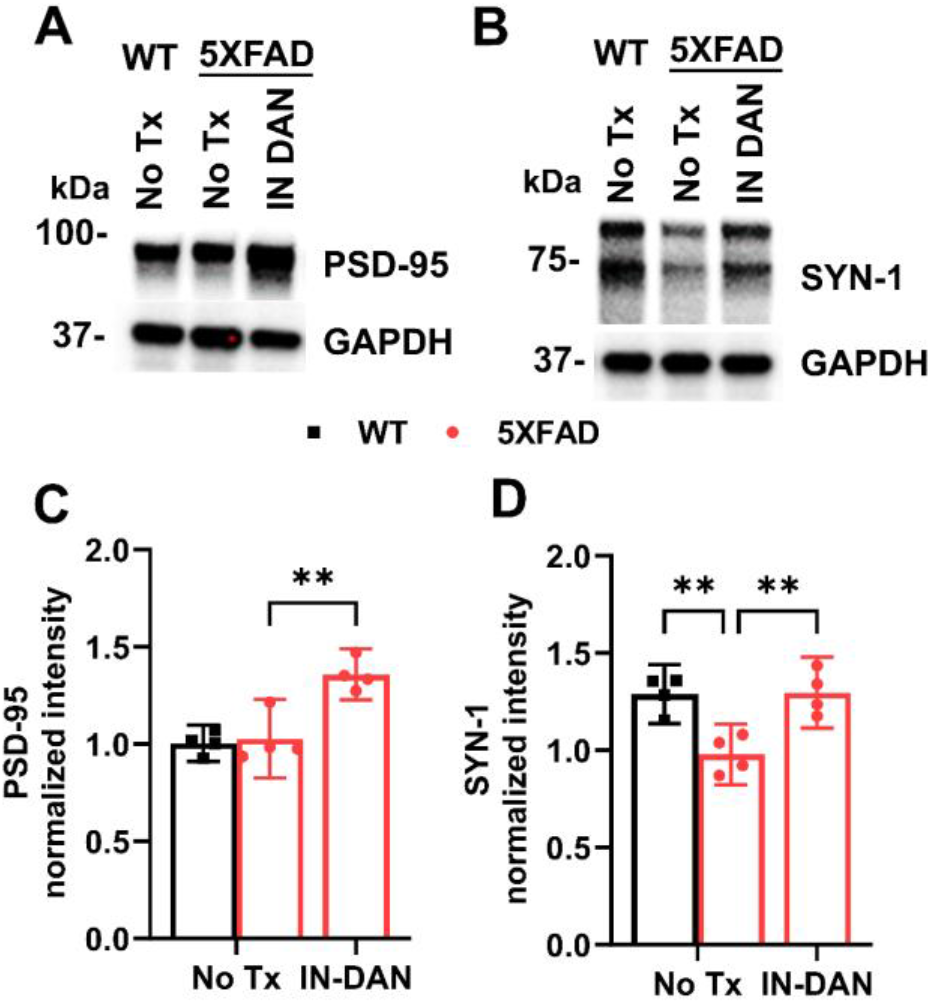
Dantrolene inhibits synapse loss in aged 5XFAD mice brains. Wild type (WT) or 5XFAD mice at 9 months old were treated with intranasal dantrolene nanoparticles (IN-DAN, 5 mg/kg) nanoparticles, daily, Monday to Friday for 12 consecutive weeks. Mice brains were then harvested at the age of 13 months old. Western blot determined the PSD-95 (**A, C**) and Synapsin-1 (**B, D**) synapse protein levels. GAPDH as the housekeeping protein. Data are means with 95% confidence interval (Cl) from 4 separate mice brains (N=4) and were analyzed using two-way ANOVA followed by Tukey’s multiple comparison test. ^**^, P<0.01.

### Chronic administration of intranasal dantrolene nanoparticles did not cause organs toxicity or side effects

Dantrolene is a muscle relaxants and chronic use may cause multiple side effects including muscle weakness or organ toxicity such as hepatotoxicity^12^. No significant changes on body weight were detected during intranasal dantrolene nanoparticles treatment from 9 to 12 months old (Supplemental Figure 1). We used various ELISA kits to measure the blood biomarkers for major organ function, including liver (ALT), kidney (Creatinine) and thyroid (TSH). Consistent with our previous study^11^, intranasal dantrolene nanoparticles treatments for 12 weeks did not cause significant changes of liver (Supplemental Fig 2A), kidney (Supplemental Fig 2B) and thyroid functions (Supplemental Fig. 2C).

## Discussion

Intranasal administration of drugs, especially in nanoparticle formulation, significantly promotes bypass of the blood brain barrier (BBB) and penetrates the CNS, with reduces peripheral toxicity^21^. Our recent pioneer studies indicated that intranasal dantrolene nanoparticles achieved higher therapeutic efficacy in the brain compared to oral and subcutaneous administration^11, 17, 18^. Intranasal dantrolene nanoparticles significantly ameliorated memory loss in adult 5XFAD mice as a disease-modifying drug,^11^ and inflammation-related depression and anxiety behaviors in 5XFAD mice^22^. Although several studies demonstrated inhibitory effects of dantrolene on amyloid load^14, 15, 23,^ intranasal dantrolene nanoparticles did not affect amyloid load in 5XFAD mice, despite its effective amelioration of memory loss^11^. Increasing studies suggest ryanodine receptor overactivation and associated Ca^2+^ dysregulation is an upstream critical pathology leading to multiple downstream pathologies including mitochondria dysfunction, oxidative stresses, pathological inflammation, and neuron damage by pyroptosis^1^. Eventually, these pathologies result in both cognitive dysfunction and psychiatric disorders in AD^24^. This study clearly demonstrated that inhibition of RyRs overactivation by the receptor antagonist dantrolene robustly suppressed pathological inflammation and programmed cell death by inflammatory pyroptosis. This study further strengthens the indication that upstream RyRs overactivation and Ca^2+^ dysregulation and associate downstream inflammation and pyroptosis is critical pathology in AD and could be therapeutic target for effective treatments.

It has been reported that 5XFAD mice has significantly increased the NLRP3 inflammasome activation and related microgliosis and neuroinflammation^25^. In consistence, we demonstrated significant increase of NlPR3 proteins and associated pathological increase of cytotoxic proteins (IL-1β, IL-18, IL-6, TNF-a) and decrease of cytoprotective cytokines (IL-10) in 5XFAD aged mice brains, which were robustly inhibited by intranasal dantrolene nanoparticles. Moreover, intranasal dantrolene nanoparticles inhibit pathological inflammation by robustly suppressing microgliosis and astrogliosis in 5XFAD mice brains.

Our previous study indicated that dantrolene significantly inhibits synaptic proteins loss and impaired synaptogenesis in prefrontal cortical neurons derived from AD patients’ iPSCs than in that of healthy human beings^5^. It is well documented that synaptic degeneration is characterized by progressive cognitive dysfunction in AD pathogenesis^26^. Our results demonstrated that PSD-95 and synapsin-1 proteins level in dantrolene treated 5xFAD mice brains are significantly higher than the WT controls, suggesting that intranasal dantrolene nanoparticles may promote neurogenesis and synaptogenesis and keep or improve synapse function in brain regions, which need further investigation.

Since AD is a chronic disease, they need long-term drug treatments. It is important that the drugs have limited side effects or organs toxicity with chronic treatment. The major advantage using intranasal dantrolene nanoparticles administration in commonly used oral or subcutaneous approach is its significantly increased dantrolene brain/blood concentration ratio^11, 17^ which promotes its CNS therapeutic effects but minimize its side effects or organ toxicity^11^. Our previous study demonstrated no side effects on nose structure and smell function^11, 27^ liver structure^11^ and function or muscle function^11, 14, 28^ after up to 10 months chronic treatment in adult 5XFAD mice^11^. This study further demonstrated that intranasal dantrolene nanoparticles up to 12 weeks of treatment did not affect liver, kidney, thyroid functions in 5XFAD mice, or even trend to protect kidney function. These results consistently suggest that chronic intranasal dantrolene nanoparticles have minimal side effects or organs toxicity with chronic use so that it could be a suitable candidate for chronic treatment of both dementia and depression in AD patients.

This study has following limitations: 1). The sample size in each experimental group is small. However, dantrolene demonstrated statistically significant neuroprotection in 5XFAD mice brains. 2). We did not have the group of intranasal dantrolene treatment in wild type mice. However, our previous study showed that intranasal dantrolene nanoparticles did not change cognitive function in adult wild type mice^11^. 3). We were unable to measure the changes of cytosol versus mitochondria Ca2+ levels in the brain tissue, but with dantrolene inhibiting the pathological increase of both RyRs and InsP_3_R-1 protein levels, an indirect determination was made that dantrolene may inhibit upstream Ca2+ dysregulation. 4). We did not measure the contents of reactive oxygen species (ROS) concentrations in brains but used the changes of oxidized lipid membrane proteins as indirect indication of oxidative stresses in the brain.

## Conclusion

In conclusion, intranasal dantrolene nanoparticles significantly inhibited pathological increased RyR-2 and InsP_3_R-1 Ca^2+^ channels proteins levels, oxidative stresses on lipid membrane proteins, pathological inflammation, programmed cell death by pyroptosis and synaptic protein loss, without side effects or organs toxicity on liver, kidney or thyroid with chronic use. This study provides a new mechanism of dantrolene neuroprotection in AD.

## Acknowledgements

This work was supported by grants to HW from the National Institute on Aging (R01AG061447) and NIA R01 Supplemental (3R01AG061447-03S1). The research was performed in the lab of Dr. Huafeng Wei and should be attributed to the Department of Anesthesiology, University of Pennsylvania. We appreciate technical support from Yutong Yi, an undergraduate student at UPENN.

## Author contributions

H.W. conceived and designed the study. P.B, W.Z, R.S., K.K, Y.W, G.L. conducted, acquired and the data, P.B, W.Z, R.S., K.K., Y.W., G.L. M.E. and H.W. analyzed data and contributed to the manuscript preparation. All the authors reviewed and approved the final manuscript.

## Data Availability

Data is available upon request.

---

**Supplemental Fig. 1.**
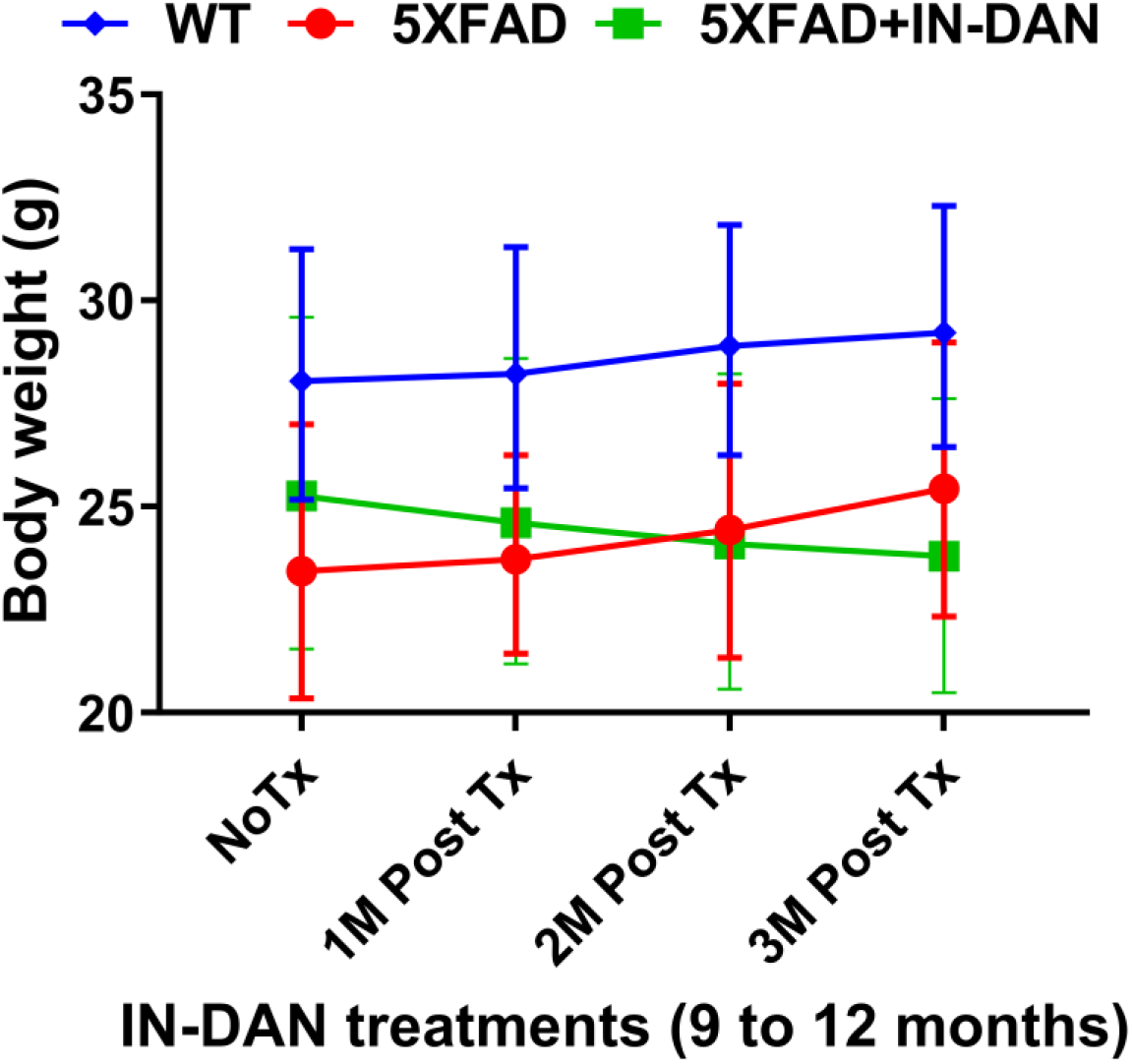
Chronic intranasal dantrolene nanoparticle administration did not affect body weight. Wild type (WT) or 5XFAD mice at 9 months old were treated with intranasal dantrolene nanoparticles (IN-DAN, 5 mg/kg) nanoparticles, daily, Monday to Friday for 12 consecutive weeks. Body weight was determined monthly from initiation of IN dantrolene treatment at 9 months old till the end of all behavioral tests at 12 months old. Data are means with 95% confidence interval (Cl) from 4-6 separate mice and were analyzed using two-way ANOVA.

**Supplemental Fig. 2.**
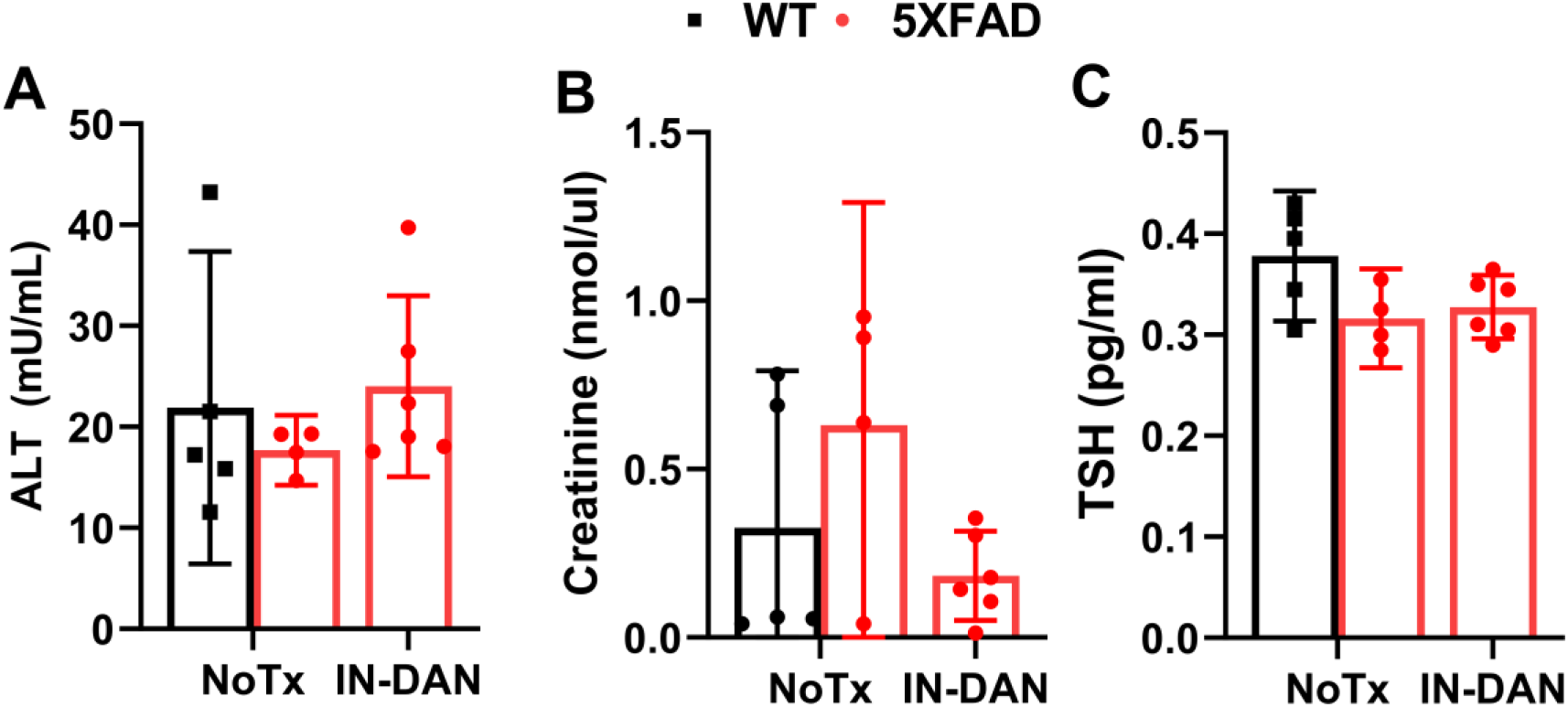
Chronic intranasal dantrolene nanoparticles administration did not cause organs toxicity. Wild type (WT) or 5XFAD mice at 9 months old were treated with intranasal (IN) dantrolene (DAN, 5 mg/kg) nanoparticles, daily, Monday to Friday for 12 consecutive weeks. Mice bloods were then harvested at the age 13 months old. Blood biomarkers for liver (ALT: Alanine Aminotransferase, **A)**, kidney (Creatinine, **B)**, and thyroid (TSH: Thyroid-stimulating hormone, **C)** function were determined using various ELISA kits. Data are means with 95% confidence interval (Cl) from 4-6 separate mice and were analyzed using two-way ANOVA.

